# Functional diversification of the BBSome: Insights from honey bee trypanosomatid parasite, *Lotmaria passim*

**DOI:** 10.1101/2024.10.10.617529

**Authors:** Xuye Yuan, Tatsuhiko Kadowaki

## Abstract

The BBSome is an octameric protein complex crucial for ciliary transport, though it also participates in multiple other cellular processes. These diverse functions may result from the co-option of its ancestral roles. Studying the BBSome in flagellated protists can provide insights into these ancestral functions and their subsequent adaptations. Here, we examine the function of the BBSome in *Lotmaria passim*, a monoxenous trypanosomatid parasite infecting honey bee. Parasites deficient in LpBBS2 are smaller and less motile compared to wild-type. Although intraflagellar transport of a marker membrane protein is only mildly impaired, its association with lipid rafts is significantly disrupted in the mutants. This suggests that the BBSome is essential for maintaining lipid raft integrity in *L. passim*. Transcriptomic comparisons between wild-type and LpBBS2-deficient parasites reveal that the BBSome may also influence processes related to metabolism, membrane localization of specific proteins, DNA repair, microtubules, and mitochondria. In contrast to *Leishmania mexicana*, the BBSome in *L. passim* is crucial for efficient infection of the honey bee gut, demonstrating that its cellular functions vary between related trypanosomatid species. The BBSome is likely an adaptor that links multiple proteins in a species-specific manner under various cellular contexts.

## Introduction

The assembly of cilia specific proteins requires three critical processes: the trafficking of proteins to the cilium, selective passage through the transition zone (Reiter, Blacque et al. 2012), and intraflagellar transport (IFT) (Kozminski, Johnson et al. 1993). The BBSome, an octameric complex of BBS1, BBS2, BBS4, BBS5, BBS7, BBS8, BBS9, and BBS18 (Chou, Apelt et al. 2019, Klink, Gatsogiannis et al. 2020, Singh, Gui et al. 2020, Yang, Bahl et al. 2020), was originally suggested to function for ciliary transmembrane protein import (Berbari, Lewis et al. 2008, Jin, White et al. 2010). The BBSome is enriched at the transition zone (Dean, Moreira-Leite et al. 2016) and moves bidirectionally during IFT associated with the IFT-A and IFT-B complexes (Williams, McIntyre et al. 2014). Accumulation of the transmembrane proteins in cilium by the loss of BBSome led to the hypothesis that the BBSome promotes the export of ciliary transmembrane proteins (Nachury 2018, Lechtreck 2022), while IFT-A stimulates their entry (Hirano, Katoh et al. 2017). Indeed, the BBSome was involved in the exit of phospholipase D, SSTR3, and Smoothened from the cilium (Lechtreck, Brown et al. 2013, Ye, Nager et al. 2018). In addition to the ciliary functions, BBSome has several other intracellular functions including regulating intracellular vesicular traffick (Guo, Cui et al. 2016), cell cytoskeleton dynamics (Hernandez-Hernandez, Pravincumar et al. 2013), gene expression (Gascue, Tan et al. 2012), proteasome-dependent protein degradation (Liu, Tsai et al. 2014), and mitochondrial homeostasis (Guo, Merrill et al. 2023). Many phenotypes associated with BBSome deficiency could be explained by ciliary defects but some may also result from the non-ciliary functions.

Despite extensive studies on the BBSome in mammals, there are few reports on its role in trypanosomatid parasites. *Leishmania major* lacking BBS1 shows no defects in growth, flagellar assembly, motility, or differentiation *in vitro* but fails to establish infection in mice (Price, Paape et al. 2013). Similarly, in *Trypanosoma brucei*, the BBSome is dispensable for flagellar assembly, motility, bulk endocytosis, and cell viability but required for virulence. These findings suggest that the BBSome facilitates endocytic sorting of specific membrane proteins at the ciliary base through interactions with clathrin and ubiquitin (Langousis, Shimogawa et al. 2016). However, its role in ciliary transport in these parasites remains unclear.

Given the conservation of BBSome subunits in flagellated protists, studying its function in different organisms can provide insight into its ancestral roles. In this study, we focus on *Lotmaria passim*, a monoxenous trypanosomatid that infects honey bee hindguts globally (Ravoet, Schwarz et al. 2015). This parasite negatively impacts honey bee health (Liu, Lei et al. 2020). Our investigation reveals that while LpBBS1 is essential for viability, LpBBS2 is necessary for proper morphogenesis, motility, flagellar transport, lipid raft integrity, and successful honey bee infection. Diverse functions of BBSome for various cellular processes in flagellated protists will be discussed.

## Results

### Intracellular Localization of LpBBS1 and LpBBS2

To explore the role of the BBSome complex in *L. passim*, we examined the intracellular localization of LpBBS1 and LpBBS2 by tagging them with triple c-Myc epitopes and expressing them using episomal vectors. As shown in Figure 1, both LpBBS1 and LpBBS2 are distributed throughout the cell body, with significant enrichment at the anterior end. The endogenous LpBBS1, tagged with triple c-Myc epitopes and degron (miniIAA7), exhibited the same localization pattern, indicating that this is not an artifact of ectopic protein expression. This localization was absent in wild-type *L. passim*. In contrast, a subunit of IFT-B complex, LpIFT88-GFP was confined to the base of the flagellum. Previous studies have shown that the BBSome localizes to the flagellar pocket membrane in *T. brucei* and that BBS1 is present throughout the cell body in *L. major* (Price, Paape et al. 2013, Langousis, Shimogawa et al. 2016). These results suggest that the BBSome likely performs multiple roles in *L. passim*.

**Figure 1.**
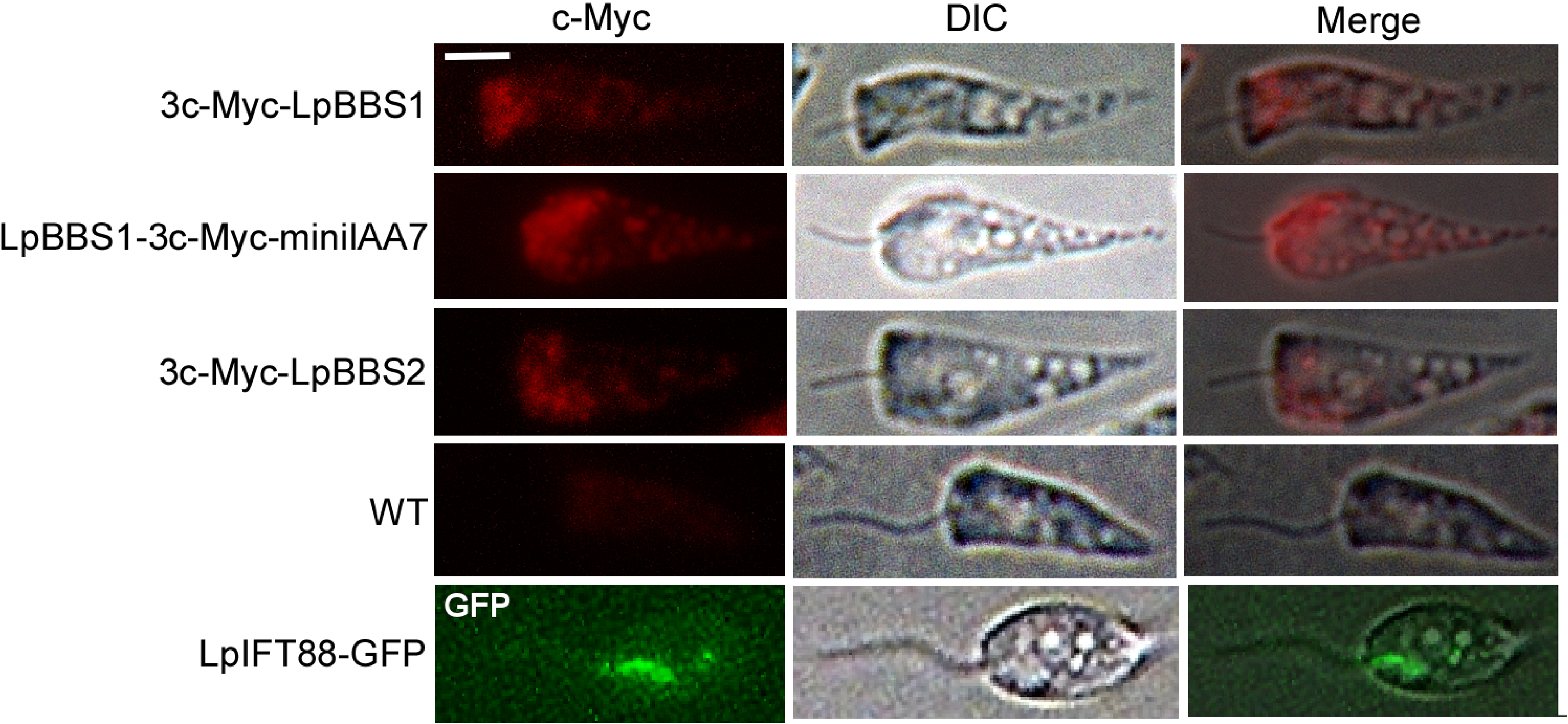
Cellular localizations of LpBBS1, LpBBS2, and LpIFT88-GFP. *Lotmaria passim* expressing 3c-Myc-LpBBS1, LpBBS1-3c-Myc-miniIAA7, or 3c-Myc-LpBBS2, along with wild-type (WT) controls, were analyzed via immunofluorescence (c-Myc) and differential interference contrast (DIC) microscopy. LpIFT88-GFP was directly visualized using fluorescence light (GFP). Merged images are provided. The anterior end of each parasite is oriented to the left. Scale bar: 2 μm.

### Application of Auxin-Inducible Degron (AID) System to Study LpBBS1 Function

We attempted to delete the *LpBBS1* gene using CRISPR but were unable to generate homozygous knock-out clones, suggesting that LpBBS1 is essential for *L. passim* viability. To further study its function, we tested the applicability of the auxin-inducible degron (AID) system (Li, Prasanna et al. 2019, Yesbolatova, Saito et al. 2020) in *L. passim*. We created parasites expressing a miniIAA7-tagged GFP and *Arabidopsis thaliana* AFB2 (AtAFB2) on episomal vectors. Upon the addition of IAA (auxin), the miniIAA7-tagged GFP was rapidly degraded (Fig. 2A), confirming the functionality of the AID system in *L. passim*.

**Figure 2.**
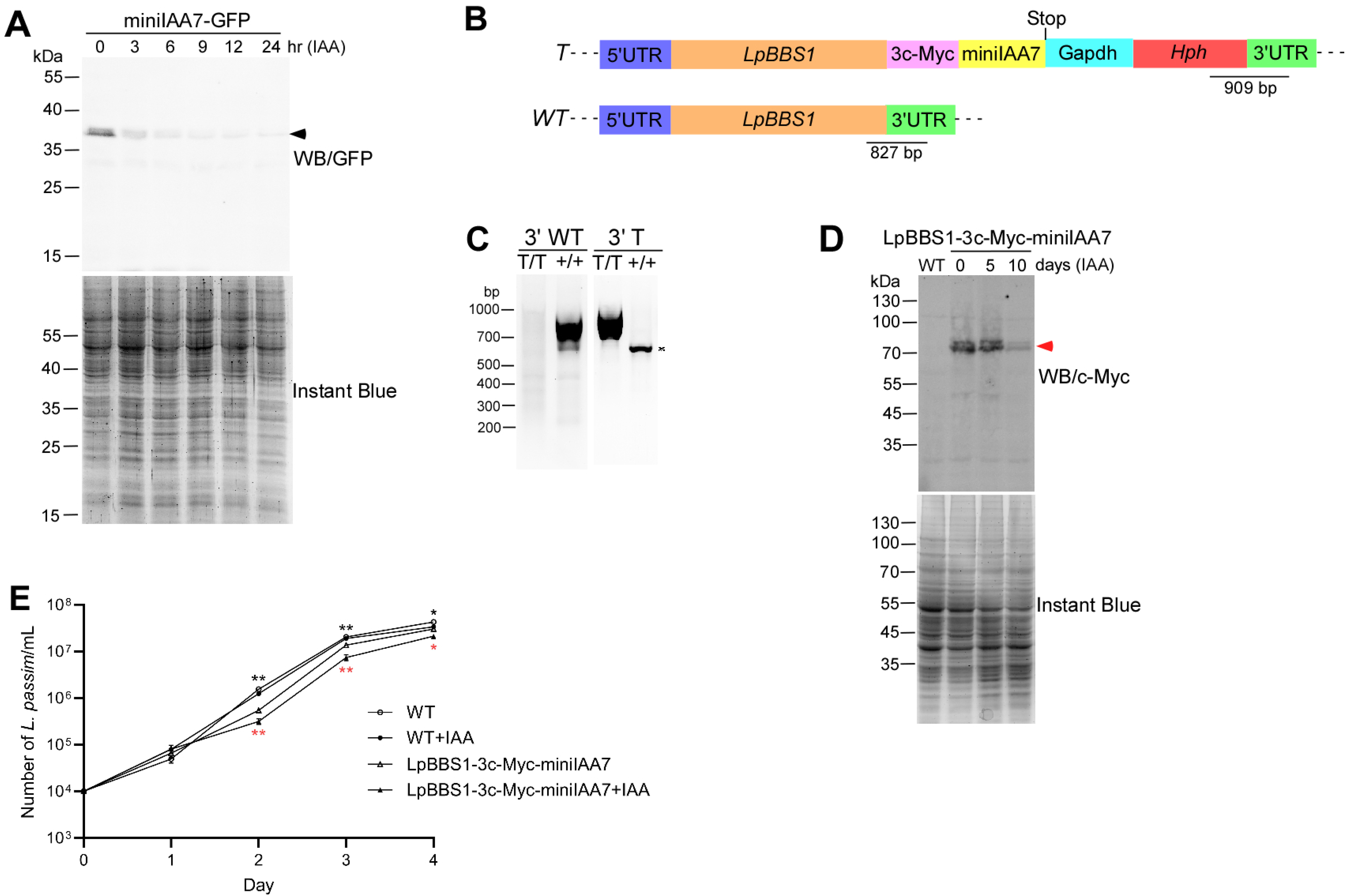
AID-mediated functional analysis of LpBBS1. (A) *L. passim* expressing degron-tagged GFP (miniIAA7-GFP) and AtAFB2 was treated with IAA for 0–24 hours. Cell lysates were analyzed by 12% SDS-PAGE, followed by western blot with anti-GFP antibody (WB/GFP) and Instant Blue staining. The miniIAA7-GFP band is marked by an arrowhead. Protein marker molecular weights (kDa) are indicated on the left. (B) Schematic representation of wild-type (*WT*) and tagged (*T*) alleles of LpBBS1, created via CRISPR/Cas9-induced homology-directed repair. The 5’ and 3’ untranslated regions (UTRs), open reading frame (ORF), 3c-Myc epitopes, miniIAA7, *Trypanosoma cruzi* Gapdh terminator, and hygromycin resistance gene (Hph) are color-coded. The stop codon is indicated. Expected PCR product sizes for detecting *WT* and *T* alleles (not to scale) are also shown. (C) Genomic DNA from WT (+/+) and homozygous tagged *LpBBS1* clones (T/T) was analyzed by PCR to detect 3’ *WT* and 3’ *T* alleles. Molecular weight marker sizes are shown on the left. A non-specific PCR band is marked with an asterisk. (D) Cell lysates from LpBBS1-3c-Myc-miniIAA7 parasites expressing AtAFB2, treated with IAA for 0–10 days, and untreated WT parasites were analyzed by 8% SDS-PAGE. LpBBS1-3c-Myc-miniIAA7 protein (red arrowhead) was detected via western blot using anti-c-Myc antibody (WB/c-Myc), with total proteins visualized by Instant Blue staining. (E) Growth rates of WT (white circles), WT with IAA (+IAA, black circles), LpBBS1-3c-Myc-miniIAA7 (clone E9, white triangles), and LpBBS1-3c-Myc-miniIAA7 with IAA (+IAA, black triangles) were monitored in culture for 4 days at 30°C (biological replicates, n = 3). Growth differences between WT and LpBBS1-3c-Myc-miniIAA7 without IAA (black asterisks, day 2; *P* < 0.002, day 3; *P* < 0.003, day 4; *P* < 0.02) and between treated and untreated LpBBS1-3c-Myc-miniIAA7 (red asterisks, day 2; *P* < 0.003, day 3; *P* < 0.002, day 4; *P* < 0.02) were statistically significant (two-tailed Welch’s t-test).

To tag the endogenous LpBBS1 with triple c-Myc epitopes and miniIAA7, we constructed a homology-directed repair (HDR) plasmid DNA made with 3’ORF of *LpBBS1*, triple c-Myc epitopes, miniIAA7, *Trypanosoma cruzi Gapdh* terminator sequence derived from pTrex-n-eGFP (Peng, Kurup et al. 2014), hygromycin-resistant gene (*Hph*), and truncated 3’UTR of *LpBBS1* (Fig. 2B). This construct was electroporated to the parasites expressing Cas9 and gRNA targeted at 29 bp downstream of the stop codon of *LpBBS1*. Successful integration of this construct into the parasite genome was confirmed by PCR (Fig. 2C), creating homozygous tagged clones (LpBBS1-3c-Myc-miniIAA7). Western blot analysis confirmed the expression of the tagged LpBBS1.

However, depletion of the tagged LpBBS1 upon IAA treatment was slow, requiring more than five days, and the protein remained detectable even after 10 days (Fig. 2D). We were able to maintain the LpBBS1-3c-Myc-miniIAA7 clone in the continuous presence of IAA, but its growth was slower compared to the untreated one. Moreover, the LpBBS1-3c-Myc-miniIAA7 clone proliferated more slowly than the wild type, even without IAA, at 30℃ (Fig. 2E). These results indicate that LpBBS1 is crucial for normal growth in *L. passim* and that tagging with triple c-Myc epitopes and miniIAA7 impairs its function under culture conditions. Disruption of *BBS1* genes in *L. major* and *T. brucei* did not affect the normal growth (Price, Paape et al. 2013, Langousis, Shimogawa et al. 2016), suggesting that LpBBS1 may carry out the functions that are absent in the above trypanosomatid parasites.

### Role of LpBBS2 in Proliferation, Morphology, and Motility

Given the challenges in depleting LpBBS1, we focused on LpBBS2. Using CRISPR, we successfully disrupted the *LpBBS2* gene, replacing its partial open reading frame (1-1271) with *Hph* gene (Fig. 3A). PCR confirmed the loss of wild-type alleles at both the 5’ and 3’ ends of the gene (Fig. 3B), and the absence of *LpBBS2* mRNA was verified (Fig. 3C). Unlike LpBBS1, LpBBS2 was not essential for *L. passim* viability under culture conditions. Growth analysis revealed that while the *LpBBS2* mutant grew normally at 30°C (Fig. 4A), it exhibited dramatically reduced growth at 21°C (Fig. 4B). In the early stationary phase (three days after culture initiation at 30℃), wild-type parasites displayed active movement with extended flagella. However, LpBBS2-deficient parasites had shorter flagella, smaller cell bodies, and reduced motility compared to wild-type parasites (Fig. 4C-F and Supplementary Videos 1 and 2). These findings indicate that LpBBS2 plays a critical role in the morphology and motility of *L. passim*.

**Figure 3.**
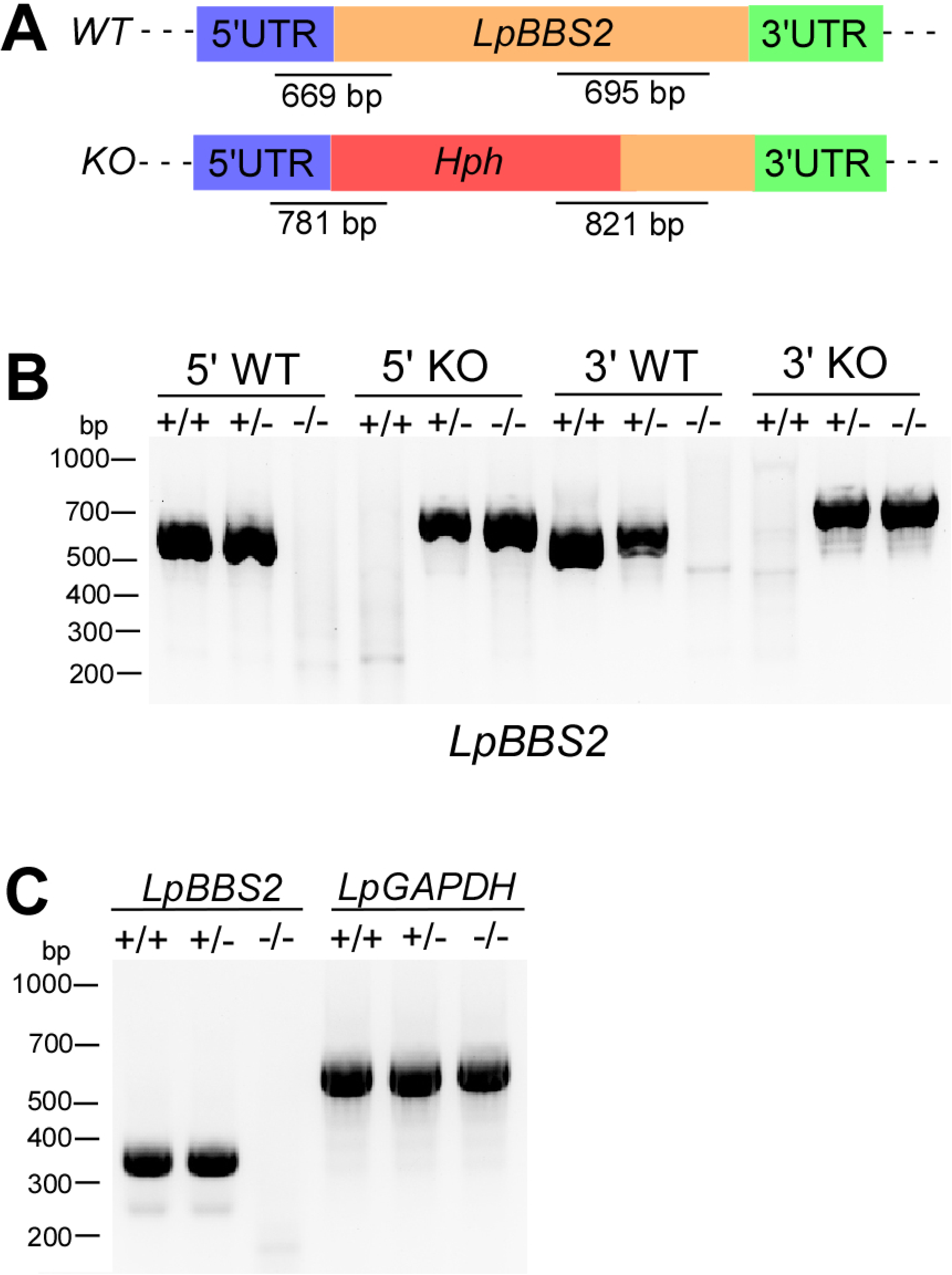
CRISPR-mediated deletion of *LpBBS2*. (A) Schematic representation of *WT* and deleted (*KO*) alleles of *LpBBS2* generated via CRISPR/Cas9-induced homology-directed repair. 5’ and 3’ UTRs, ORF, and *Hph* are color-coded. Expected PCR product sizes for detecting *WT* and *KO* alleles (not to scale) are shown. (B) Genomic DNA from WT (+/+), heterozygous (+/-), and homozygous (-/-) *LpBBS2* mutant parasites was analyzed by PCR for *5’ WT*, *5’ KO*, *3’ WT*, and *3’ KO* alleles. Molecular weight markers are shown on the left. (C) RT-PCR detection of *LpBBS2* and *LpGAPDH* mRNAs in *LpBBS2* heterozygous and homozygous mutants, along with WT parasites, using a forward primer targeting the *L. passim* splice leader sequence. Molecular weight markers are shown on the left.

**Figure 4.**
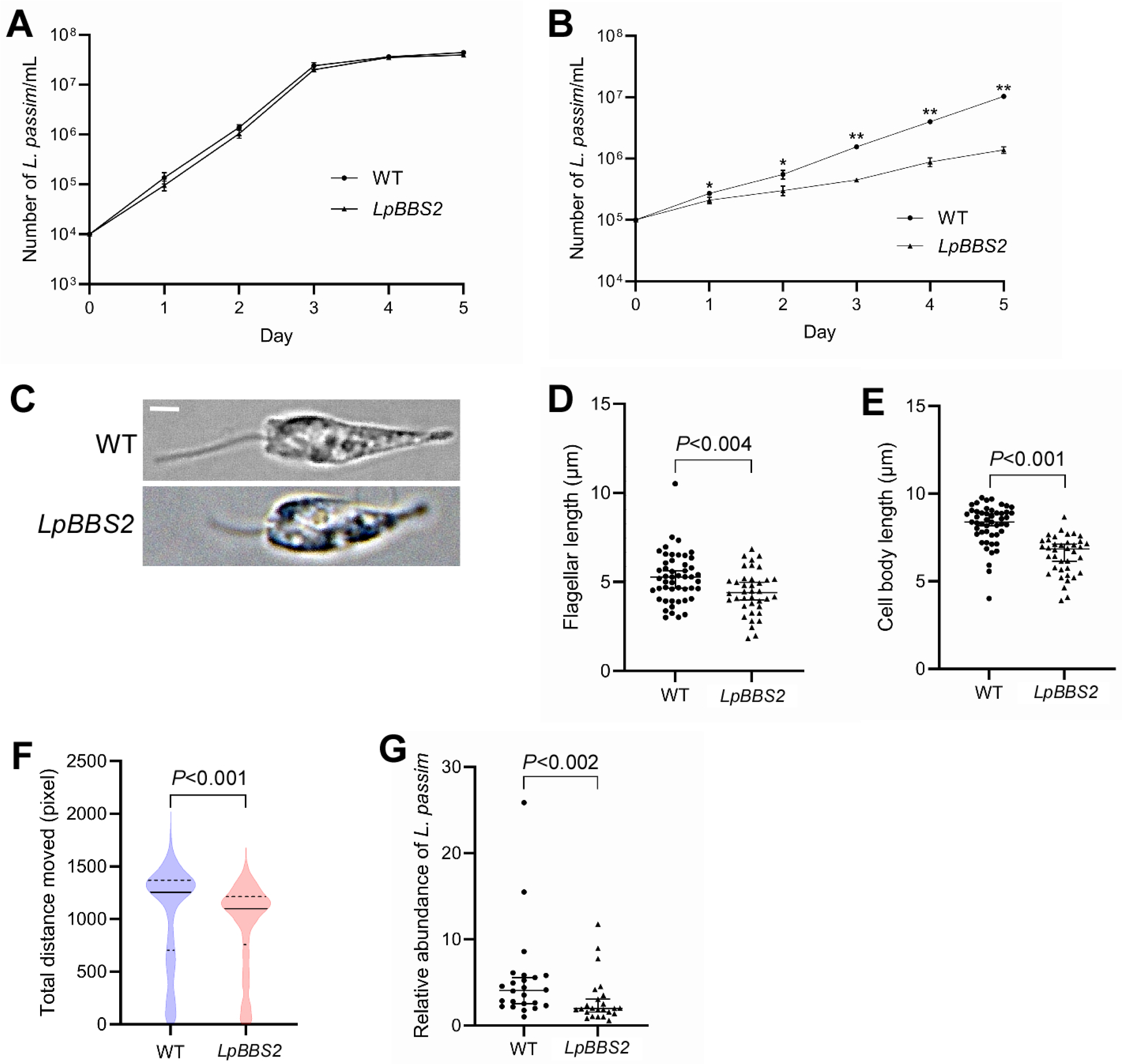
Phenotypes of LpBBS2-deficient mutants in culture and honey bees. (A) Growth rates of WT (circles) and *LpBBS2* homozygous mutants (clone D8, triangles) were measured at 30°C over 5 days (biological replicates, n = 3). (B) Growth rates of WT and *LpBBS2* homozygous mutants were measured at 21°C over 5 days (biological replicates, n = 3). Statistically significant differences in growth rates between WT and *LpBBS2* mutants across days 1–5 are indicated (asterisks, day 1; *P* < 0.04, day 2; *P* < 0.03, day 3; *P* < 0.002, day 4; *P* < 0.00002, day 5; *P* < 0.0004). (C) Morphology of WT and *LpBBS2* mutants. Scale bar: 2 μm. (D) Flagellar length comparisons between WT (n = 49) and *LpBBS2* mutants (n = 38). Statistical analysis was performed using the Brunner-Munzel test. (E) Cell body length comparisons between WT (n = 49) and *LpBBS2* mutants (n = 38). (F) Motility (total distance moved in 1 minute) of WT (n = 1,415) and *LpBBS2* mutants (n = 1,178) is shown via violin plot. Median, first, and third quartiles are indicated by solid and dashed lines, respectively. (G) Relative abundance of *L. passim* in individual honey bees (n = 24) 14 days post-infection, comparing WT and *LpBBS2* mutants. Data is normalized to one sample infected with WT *L. passim* as 1, and the median with 95% CI is presented.

### LpBBS2 and Lipid Raft Integrity

We previously reported that the flagellar calcium binding proteins (FCaBPs) localize in the flagellum (LpFCaBP1) and entire body (LpFCaBP2) membrane of *L. passim* through the specific N-terminal sorting sequences modified by myristoylation and palmitoylation (Yuan and Kadowaki 2024). Given that LpFCaBP1 interacts with the BBSome, we examined the localization of LpFCaBP1N16-GFP and LpFCaBP2N16-GFP (GFP fused with N-terminal 16 amino acids of LpFCaBP1 and LpFCaBP2) in wild-type and *LpBBS2* mutants. In wild-type parasites, LpFCaBP1N16-GFP was enriched in the flagellum, but in the LpBBS2 mutant, a significant amount was mislocalized to the cell body (Fig. 5A and B). LpFCaBP2N16-GFP remained in the cell body membrane of both wild-type and LpBBS2 mutants. FCaBP was shown to associate with lipid rafts in the flagellum of *T. brucei* and *T. cruzi* (Tyler, Fridberg et al. 2009, Maric, McGwire et al. 2011). This is consistent with that palmitoylation and myristoylation promote the association of protein with lipid rafts (Aicart-Ramos, Valero et al. 2011). We thus tested lipid raft association of LpFCaBP1N16-GFP and LpFCaBP2N16-GFP in wild-type and *LpBBS2-*deficient parasites. Both proteins were present in the pellet (insoluble) fraction when wild-type and *LpBBS2* mutant were extracted with 1% TX-100 at 4℃ but shifted to the soluble fraction at 37℃ (Fig. 5C). These results demonstrate that LpFCaBP1N16-GFP and LpFCaBP2N16-GFP associate with lipid rafts in both wild-type and *LpBBS2* mutant. When the parasites were extracted at 20℃, most of both proteins were present in the pellet fraction for wild-type but in the soluble fraction for *LpBBS2* mutant (Fig. 5C and D). These results suggest their raft association was weakened in the *LpBBS2* mutant at 20°C. These findings suggest that LpBBS2 is involved in maintaining lipid raft integrity.

**Figure 5.**
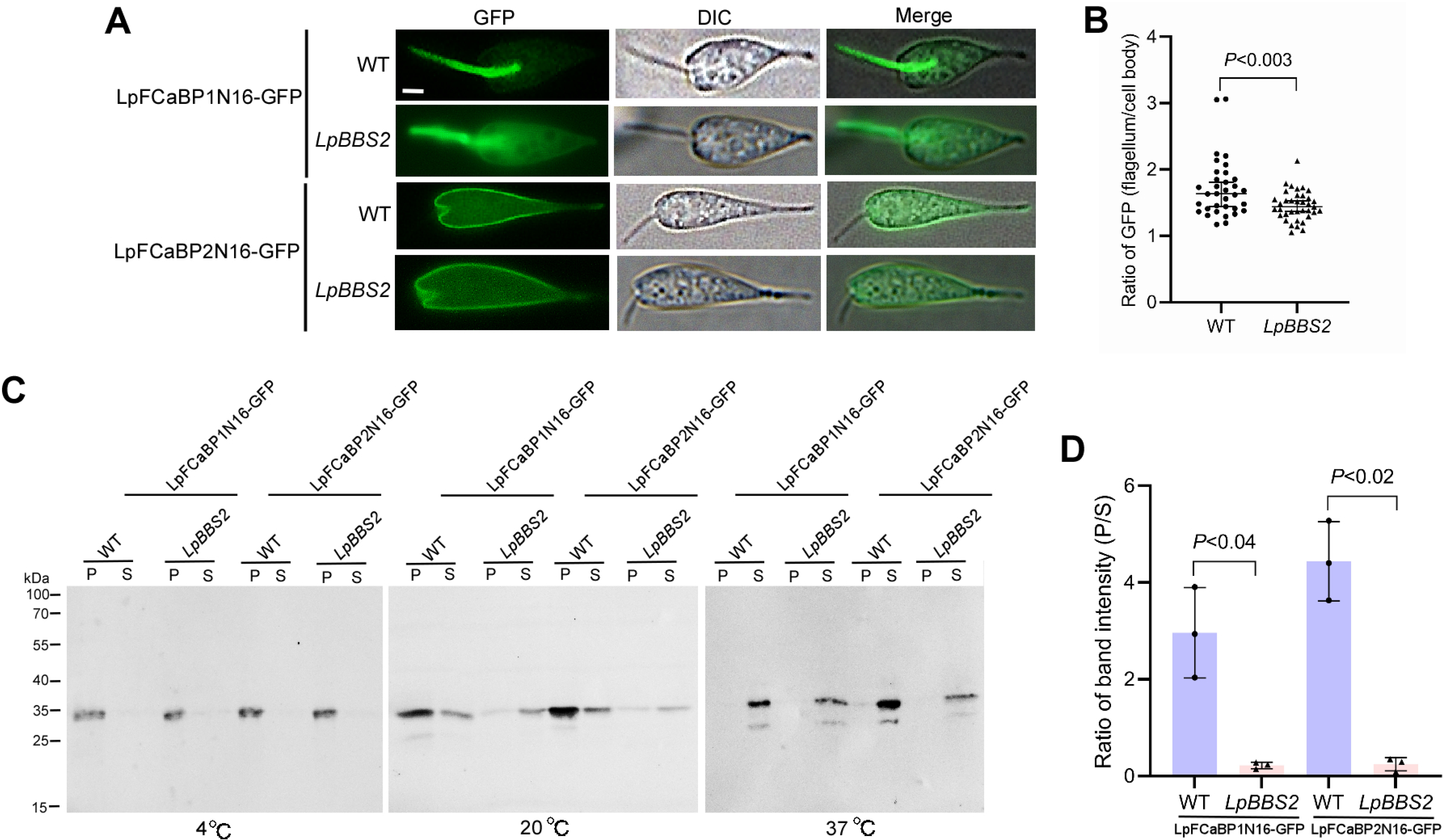
Localization and lipid raft association of LpFCaBP1N16-GFP and LpFCaBP2N16-GFP in WT and LpBBS2-deficient parasites. (A) WT and *LpBBS2* mutant *L. passim* expressing LpFCaBP1N16-GFP or LpFCaBP2N16-GFP were observed under DIC and fluorescence microscopy. Merged images are shown. Scale bar: 2 μm. (B) Comparison of GFP fluorescence ratios (flagellum to cell body) between WT (n = 34) and *LpBBS2* mutants (n = 37). Statistical analysis was conducted using the Brunner-Munzel test. (C) WT and *LpBBS2* mutant parasites expressing LpFCaBP1N16-GFP or LpFCaBP2N16-GFP were solubilized in 1% Triton X-100 at different temperatures, and the soluble (S) and pellet (P) fractions were analyzed by western blot using anti-GFP antibody. Molecular weights of protein markers are indicated on the left. (D) GFP band intensity ratio (P/S) from parasites solubilized at 20°C is compared between WT and *LpBBS2* mutants (biological replicates, n = 3). Statistical analysis was performed using a two-tailed Welch’s t-test.

### Transcriptomic Changes in *LpBBS2* Mutants

To uncover the functions of the BBSome from a more comprehensive perspective of the entire cell, we performed RNA-seq to investigate the broader effects of *LpBBS2* deletion on the *L. passim* transcriptome. Principal component analysis revealed distinct transcriptomes for LpBBS2 mutant clones (D8 and G1) compared to wild-type (Supplementary Figure 1A). A scatter plot analysis showed differential expression of over 2,800 genes between the mutants and wild-type. Expression of only 40 genes was significantly different between the clones D8 and G1, indicating their similarity (Supplementary Figure 1B-D). GO-term and KEGG pathway enrichment analyses suggested that upregulated genes in *LpBBS2* mutant were associated with metabolism, cell cycle, cellular response to extracellular stimulus, DNA repair, cytoskeleton, mitochondria, and nucleus. While downregulated genes were linked to ribosome biogenesis and proteasome (Tables 1-3). Among genes related to IFT, we found that dynein light and heavy chain genes are up-regulated in *LpBBS2* mutant (Supplementary Table 1). These transcriptomic changes suggest that LpBBS2 plays a broad regulatory role in cellular processes.

**Table 1.** Representative GO-terms enriched with genes up-regulated in *LpBBS2* mutants.

| ONTOLOGY | ID | Description | Adjusted *P*-value |
| --- | --- | --- | --- |
| BP | GO:0051493 | regulation of cytoskeleton organization | 0.024512417 |
| BP | GO:1901360 | organic cyclic compound metabolic process | 0.030397066 |
| BP | GO:0033043 | regulation of organelle organization | 0.031204177 |
| BP | GO:0007346 | regulation of mitotic cell cycle | 0.031204177 |
| BP | GO:0006766 | vitamin metabolic process | 0.031204177 |
| BP | GO:0016311 | dephosphorylation | 0.046513742 |
| BP | GO:1901615 | organic hydroxy compound metabolic process | 0.046513742 |
| BP | GO:0031668 | cellular response to extracellular stimulus | 0.049831369 |
| BP | GO:0032886 | regulation of microtubule-based process | 0.049831369 |
| BP | GO:0051103 | DNA ligation involved in DNA repair | 0.049831369 |
| BP | GO:1901700 | response to oxygen-containing compound | 0.049831369 |
| CC | GO:0005739 | mitochondrion | 0.00011969 |
| CC | GO:0005634 | nucleus | 0.024513222 |
| CC | GO:0005814 | centriole | 0.045586455 |
| CC | GO:0005874 | microtubule | 0.045586455 |
| CC | GO:0090734 | site of DNA damage | 0.045586455 |

BP: Biological process, CC: Cellular compartment

**Table 2.** Representative GO-terms enriched with genes down-regulated in *LpBBS2* mutants.

| ONTOLOGY | ID | Description | Adjusted *P*-value |
| --- | --- | --- | --- |
| BP | GO:0006364 | rRNA processing | 0.000412333 |
| BP | GO:0034470 | ncRNA processing | 0.000412333 |
| BP | GO:0042254 | ribosome biogenesis | 0.000412333 |
| BP | GO:0022613 | ribonucleoprotein complex biogenesis | 0.002349856 |

**Table 3.** KEGG pathways enriched in *LpBBS2* mutants.

| Up-regulated genes |  |  |
| --- | --- | --- |
| ID | Description | Adjusted *P*-value |
| ko03029 | Mitochondrial biogenesis [BR:ko03029] | 0.000637691 |
| ko05208 | Chemical carcinogenesis - reactive oxygen species | 0.023018084 |
| ko04714 | Thermogenesis | 0.046239438 |
| Down-regulated genes |  |  |
| ko03011 | Ribosome [BR:ko03011] | 2.05511E-23 |
| ko03009 | Ribosome biogenesis [BR:ko03009] | 7.12255E-15 |
| ko03051 | Proteasome [BR:ko03051] | 0.001109833 |

### LpBBS2 is Essential for Efficient Infection in the Honey Bee Hindgut

To assess the role of LpBBS2 in *L. passim* infection, we infected honey bees with wild-type and LpBBS2-deficient parasites and measured parasite load in the hindgut after 14 days. The *LpBBS2* mutant exhibited a significantly reduced infection rate compared to wild-type (Fig. 4G), indicating that LpBBS2 is required for efficient infection of the honey bee hindgut.

## Discussion

### Application of AID in Trypanosomatid Parasites

AID has been successfully used to study gene functions in various organisms, from yeast to mammals (Li, Prasanna et al. 2019, Yesbolatova, Saito et al. 2020). In this study, we applied AID to investigate the function of LpBBS1, which is essential for the viability of *L. passim*. The degron-tagged GFP at the N-terminus was rapidly degraded upon the addition of IAA; however, this was not the case for LpBBS1, which required prolonged incubation with IAA (more than five days) to sufficiently deplete the protein (Fig. 2). The degradation rate via AID appears to depend on both the target protein and the position of the degron sequence in *L. passim*, similar to findings in other species (Li, Prasanna et al. 2019, Yesbolatova, Saito et al. 2020). Through AID, we discovered that LpBBS1 is necessary for the normal growth of *L. passim* under culture conditions. This differs from *L. major* and *T. brucei*, where BBS1 is not required for normal growth (Price, Paape et al. 2013, Langousis, Shimogawa et al. 2016). Thus, the BBSome in monoxenous trypanosomatid parasites may have additional roles not present in dixenous trypanosomatid parasites. AID is a useful approach for studying gene function in trypanosomatid parasites under culture conditions; however, its application *in vivo* could be challenging, as IAA needs to be administered to the host and remain stable.

### Cellular Functions of the BBSome in Flagellated Protists

In contrast to LpBBS1, we found that *L. passim* lacking LpBBS2 is viable. BBS2 and BBS7 subunits form the head of the BBSome by creating an asymmetric heterodimer, which connects with α-helical domains (Chou, Apelt et al. 2019, Klink, Gatsogiannis et al. 2020, Singh, Gui et al. 2020, Yang, Bahl et al. 2020). However, some species, such as fruit flies, lack the *BBS2* gene (Ewerling, Maissl et al. 2023), suggesting that the head part is not essential for the core function of the BBSome. Alternatively, the BBSome subunit may have individual function without forming the octamer. Nevertheless, we found that LpBBS2 is necessary for the normal growth of *L. passim* in culture at 21°C, but not at 30°C. Furthermore, LpBBS2-deficient parasites have shorter flagella and cell bodies compared to the wild-type, and their motility is also reduced (Fig. 4). Consistent with the role of the BBSome in the IFT of membrane proteins, the dually-acylated LpFCaBP1N16-GFP was less abundant in the flagella of *LpBBS2* mutants compared to the wild-type. Although the BBSome does not seem to be involved in the IFT of membrane proteins in *T. brucei* and *L. major* (Price, Paape et al. 2013, Langousis, Shimogawa et al. 2016), it plays an important role in other flagellated protists. In *Chlamydomonas reinhardtii*, the BBSome is required for the ciliary export of dually-acylated phospholipase D via retrograde transport (Lechtreck, Brown et al. 2013). In *Paramecium tetraurelia*, RNAi knockdown of the BBSome results in the loss of calcium-activated K^+^ channels and PKD from the cilia (Valentine, Rajendran et al. 2012). In *Euglena gracilis*, the deletion of BBS7 and BBS8 leads to the loss of flagella (Ishikawa, Nomura et al. 2022). The weak association of LpFCaBP1N16-GFP and LpFCaBP2N16-GFP with lipid rafts suggests that the lipid raft environment may be altered in *LpBBS2* mutants. Since lipid rafts in trypanosomatids are rich in 3 β–hydroxysterols and sphingolipids, similar to those in other species (Tetley 1986, Tyler, Fridberg et al. 2009, Serricchio, Schmid et al. 2015), enzymes involved in the metabolism of these lipids may be affected. Alternatively, the lipid rafts associated proteins necessary to anchor FCaBPs would be changed in the mutants. Although LpFCaBP2N16-GFP is highly enriched in the cell body membrane of *LpBBS2* mutants (Fig. 4A), we can not completely rule out the possibility that the DHHC palmitoyltransferases and N-myristoyltransferase, responsible for palmitoylation and myristoylation, are partially impaired. Protein acylation is known to be critical for the association with lipid rafts (Aicart-Ramos, Valero et al. 2011).

We believe the transcriptomic changes in *LpBBS2* mutants are likely direct or indirect consequences of compensating for the lack of this gene. Since dynein is the specific motor protein for retrograde transport along microtubules (Pazour, Wilkerson et al. 1998), the upregulation of dynein light and heavy chain mRNAs in *LpBBS2* mutants would be consistent with the role of the BBSome in IFT of membrane proteins. According to GO terms enriched with upregulated genes in *LpBBS2* mutants, the BBSome may play multiple roles related to metabolism, DNA repair, plasma membrane protein localization, mitochondria, and microtubules in *L. passim*. Indeed, many studies report these non-ciliary functions of the BBSome. For example, compromised cell migration, adhesion, and division in BBS4-deficient cells indicate a role for the BBSome in regulating cytoskeleton dynamics (Hernandez-Hernandez, Pravincumar et al. 2013). The BBSome is also required for the plasma membrane localization of proteins such as the leptin receptor (Guo, Cui et al. 2016), Notch (Leitch, Lodh et al. 2014), and the insulin receptor (Starks, Beyer et al. 2015), suggesting a role in intracellular vesicular trafficking. The BBSome was shown to interact with proteasomal subunits, and the loss of BBS4 depletes several subunits from the centrosomal proteasome, leading to the accumulation of specific proteins (Gerdes, Liu et al. 2007, Liu, Tsai et al. 2014). This role in protein degradation is relevant to our findings that genes involved in the proteasome and ribosome pathways are downregulated in *LpBBS2* mutants (Tables 2 and 3). The BBSome has also been shown to regulate mitochondrial morphology and function by modulating the phosphorylation and mitochondrial translocation of dynamin-like protein 1 (Guo, Merrill et al. 2023).

We found that LpBBS2-deficient parasites are ineffective at infecting the honey bee hindgut. Since LpFCaBPs-deficient parasites, which have short flagella and low motility, can still infect the honeybee hind gut (Yuan and Kadowaki 2024), it is unlikely that the low infectivity of the *LpBBS2* mutants is due to their morphological or motility defects. We speculate that the alteration of lipid rafts in *LpBBS2* mutants may be responsible for the ineffective infection. Lipid rafts in trypanosomatids appear to play roles in signal transduction, virulence factor function, and endocytosis/exocytosis (Goldston, Powell et al. 2012). In the mutant, the localization and regulation of signaling molecules that may be critical for parasite adaptation to the honey bee hind gut would likely be altered (de Paulo Martins, Okura et al. 2010). For instance, the GPI-anchored variant surface glycoprotein, essential for immune evasion, is associated with lipid rafts in *T. brucei* (Denny, Field et al. 2001). This is also true for the virulence factor GP63 in *L. major* (Denny, Field et al. 2001). Furthermore, lipid rafts are implicated in endocytosis and exocytosis, which are crucial for nutrient uptake and surface protein recycling (Corrêa, Atella et al. 2007). These are vital processes for parasite survival and infection. However, the role of LpBBS2 in lipid rafts may not be shared with *Leishmania* spp, as BBS2-deficient *L. mexicana* can reach the anterior gut and survive in sand flies (Beneke, Demay et al. 2019). *L. passim* BBSome appears to have diverse cellular functions and some of them are not shared with the related trypanosomatid parasites. BBSome has ability to be an adaptor to link any of multiple proteins in a species-specific manner under various cellular contexts.

## Materials and Methods

### Cellular localization of 3c-Myc-LpBBS1, 3c-Myc-LpBBS2, and LpIFT88-GFP in *L. passim*

To express the triple c-Myc-tagged proteins 3c-Myc-LpBBS1 and 3c-Myc-LpBBS2, two oligonucleotides, 3Myc-N-5 and 3Myc-N-3, were phosphorylated and annealed, and then cloned into the XbaI and HindIII sites of the tdTomato/pTREX-b plasmid DNA (Lander, Li et al. 2015) (ADDGENE: #68709). The full open reading frames (ORFs) of the *LpBBS1* and *LpBBS2* genes were amplified using PCR with KOD-FX DNA polymerase (TOYOBO), *L. passim* genomic DNA, and primer pairs LpBBS1-5-HindIII/LpBBS1-3-ClaI and LpBBS2-5-ClaI/LpBBS2-3-ClaI, respectively. PCR products were digested with HindIII and ClaI for *LpBBS1*, and with ClaI for *LpBBS2*. The digested products were cloned into the vector mentioned above, which had been digested with the same enzymes. To construct the LpIFT88-GFP expression vector, the ORF of *LpIFT88* was amplified using primers LpIFT88-5-XbaI and LpIFT88-3-XbaI. The PCR product was digested with XbaI and cloned into the XbaI site of the pTrex-n-eGFP plasmid (ADDGENE: #62544). LpBBS1, LpBBS2, and LpIFT88 sequences are listed in Supplementary file 1.

Actively growing *L. passim* cells (4 × 10^7^) were washed twice with 5 mL PBS and resuspended in 0.4 mL Cytomix buffer (without EDTA) containing 20 mM KCl, 0.15 mM CaCl_2_, 10 mM K_2_HPO_4_, 25 mM HEPES, and 5 mM MgCl_2_ (pH 7.6). Electroporation was performed twice (1-minute interval) using 10 μg of each plasmid DNA and a Gene Pulser X cell electroporator (Bio-Rad) with a 2-mm gap cuvette, setting the voltage to 1.5 kV, capacitance to 25 μF, and resistance to infinity. The electroporated parasites were cultured in 4 mL of modified FP-FB medium (Salathé, Tognazzo et al. 2012), and blasticidin (50 μg/mL, Macklin) or G418 (200 μg/mL, Sigma-Aldrich) was added after 24 hours for selection of drug-resistant clones.

Immunofluorescence detection of 3c-Myc-LpBBS1 and 3c-Myc-LpBBS2 was performed by washing and mounting the parasites on a poly-L-lysine-coated 8-well chamber slide, fixing with 4% paraformaldehyde, permeabilizing with 0.1% Triton X-100 in PBS (PT), and blocking with PT containing 5% normal goat serum (PTG). Samples were incubated overnight at 4°C with a rabbit anti-c-Myc polyclonal antibody (1:500 dilution, Proteintech) in PTG. After five washes with PT, the samples were incubated with Alexa Fluor 555 anti-rabbit IgG (ThermoFisher) for 2 hours at room temperature, washed again, and observed under a microscope. Live *L. passim* expressing LpIFT88-GFP were washed three times with 1 mL PBS and imaged on poly-L-lysine-coated slides using a NIKON Eclipse Ni-U fluorescence microscope with a constant exposure time of 200 msec.

### Testing the AID System Using GFP in *L. passim*

To tag GFP with a degron, the miniIAA7 sequence was amplified using primers miniIAA-5-XbaI and miniIAA-3-XbaI as well as pSH-EFIRES-B-Serpin-miniIAA7-mEGFP plasmid DNA (Li, Prasanna et al. 2019) (ADDGENE: #129719) as a template, and cloned into the pTrex-n-eGFP plasmid. The complete ORF of *AtAFB2* was amplified using primers AtAFB2-5-XbaI and AtAFB2-3-HindIII as well as pSH-EFIRES-P-AtAFB2 plasmid DNA (Li, Prasanna et al. 2019) (ADDGENE: #129715) as a template, and cloned into the tdTomato/pTREX-b plasmid. *L. passim* cells were electroporated with both plasmids (10 μg each), and drug-resistant clones were selected using G418 and blasticidin.

The parasites expressing miniIAA7-GFP and AtAFB2 were cultured inoculated at 10^6^/mL in 24-well plates with 75 μg/mL auxin (IAA, Macklin) for 0-24 hours at 30°C. Cells were collected, washed with 1 mL PBS, and lysed in 100 μL SDS-PAGE sample buffer (2% SDS, 10% glycerol, 10% β-mercaptoethanol, 0.25% bromophenol blue, 50 mM Tris-HCl, pH 6.8). Samples were heated at 95°C for 5 minutes, and 20 μL was applied to two 10% SDS-PAGE gels. One gel was stained with Instant Blue (Abcam), while proteins from the other were transferred to a nitrocellulose membrane (Pall Life Sciences).

The membrane was blocked with 5% BSA in PBST (PBS with 0.1 % Tween-20), incubated with anti-GFP polyclonal antibody (1:500 dilution, Proteintech) overnight at 4 ℃. After washing five times with PBST (5 minutes each), the membrane was incubated with IRDye 680RD donkey anti-rabbit IgG (H+L) secondary antibody (1:10,000 dilution, LI-COR Biosciences) in PBST containing 5 % skim milk at room temperature for 2 hours. Following another round of washing, the membrane was visualized using ChemiDoc MP (BioRad).

### Tagging endogenous LpBBS1 with triple c-Myc epitopes and miniIAA7 by CRISPR

To tag *LpBBS1* gene, we designed the gRNA sequence targeted at 29 bp downstream of the stop codon of *LpBBS1* using a custom gRNA design tool (http://grna.ctegd.uga.edu) (Peng and Tarleton 2015). Complementary oligonucleotides (0.1 nmole each) corresponding to the sgRNA sequences (LpBBS1gRNA3’UTR48F and LpBBS1gRNA3’UTR48R) were phosphorylated by T4 polynucleotide kinase (TAKARA), annealed, and cloned into BbsI-digested pSPneogRNAH vector (Zhang and Matlashewski 2015) (ADDGENE: # 63556). We electroporated *L. passim* expressing Cas9 (Liu, Lei et al. 2019) with 10 μg of the constructed plasmid DNA and selected the transformants by blasticidin and G418 to establish parasites expressing both Cas9 and *LpBBS1* gRNA.

For the construction of donor DNA to tag the *LpBBS1* gene, we performed a fusion of two PCR products: the *T. cruzi Gapdh* terminator sequence derived from pTrex-n-eGFP (GAPDH-HindIII and GAPDH-3-Hyg) and the ORF of the *Hph* gene derived from pCsV1300 (Park, Song et al. 2013) (Hyg-5-GAPDH and Hyg-3-EcoRI). The fusion PCR products were digested and cloned into the HindIII and EcoRI sites of pBluescript II SK(+). We PCR-amplified the truncated 3’UTR of *LpBBS1* using two primers, LpBBS1-3’UTR-F-degron and LpBBS1-3’UTR-R. The PCR product was digested and cloned into the EcoRI site of the aforementioned plasmid DNA. We constructed the full-length *LpBBS1* tagged with triple c-Myc epitopes and miniIAA7 as follows. The PCR amplicon of miniIAA7, using two primers (miniIAA7-5-HindIII and miniIAA7-3-XhoI), was digested and cloned into the HindIII and XhoI sites of pBluescript II SK(+). The DNA fragment encoding triple c-Myc epitopes was then cloned into the XbaI and HindIII sites of the aforementioned plasmid DNA. By cutting the resulting plasmid DNA with XbaI and XhoI, we obtained the DNA fragment encoding triple c-Myc epitopes and miniIAA7. The complete ORF of *LpBBS1*, amplified by two primers (LpBBS1-5-XbaI and LpBBS1-3-XbaI), along with the aforementioned DNA fragment, was cloned into the XbaI and XhoI sites of pTrex-n-eGFP. Using this plasmid DNA as a template, we amplified a DNA fragment containing the *LpBBS1* ORF (703-1776), triple c-Myc epitopes, and miniIAA7 using two primers, LpBBS1-5-degron and miniIAA7-3-ClaI. The PCR product was digested and cloned into the XhoI and ClaI sites of pBluescript II SK(+) containing the *T. cruzi Gapdh* terminator, *Hph*, and truncated 3’UTR of *LpBBS1* as described above. The donor plasmid DNA (10 μg) was linearized using NotI and used for electroporation of *L. passim* expressing both Cas9 and *LpBBS1* gRNA.

After electroporation, *L. passim* resistant to blasticidin, G418, and hygromycin (150 μg/mL, Macklin) were selected, and single parasites were cloned by serial dilutions in a 96-well plate. The genotype of each clone was determined through the detection of 3’ wild-type (WT, using LpBBS1-1681F and LpBBS1-3’UTR-down) and tagged (T, using Hyg-846F and LpBBS1-3’UTR-down) alleles for *LpBBS1* by PCR. We then electroporated two independent clones (D1 and E9) with the *AtAFB2*-expressing vector, in which the blasticidin resistance gene was replaced by the bleomycin resistance gene. The *LpBBS1*-tagged *L. passim* (LpBBS1-3c-Myc-miniIAA7) expressing *AtAFB2* were selected by hygromycin and zeocin (50 μg/mL, InvivoGen).

### AID-mediated degradation of *LpBBS1* and its effect on the growth rate of *L. passim*

*LpBBS1*-3c-Myc-miniIAA7 parasites expressing AtAFB2 were cultured with IAA for 5 and 10 days at 30 °C. Then, 10⁶ cells were collected, along with untreated and WT parasites, during the mid-log growth phase. Cell lysates were prepared as described above and applied to 8% SDS-PAGE. Rabbit anti-c-Myc (1:1000 dilution) polyclonal antibody was used for the western blot. We inoculated WT and *LpBBS1*-3c-Myc-miniIAA7 parasites expressing AtAFB2 into the culture medium at 10⁴ cells/mL in a 24-well plate and cultured with or without IAA at 30 °C. The number of parasites was counted daily using a hemocytometer for 4 days. We characterized the phenotypes of the D1 and E9 clones, and their phenotypes were the same.

### Deletion of *LpBBS2* gene by CRISPR

To delete the *LpBBS2* gene, complementary oligonucleotides corresponding to the sgRNA sequences (LpBBS2gRNA1091F and LpBBS2gRNA1091R) were processed and cloned into the pSPneogRNAH vector, as described above. We then established *L. passim* expressing both Cas9 and *LpBBS2* gRNA.

For the construction of donor DNA for the *LpBBS2* gene, we performed fusion PCR of three DNA fragments: the 5’UTR of *LpBBS2* (LpBBS2 5’UTR-F and LpBBS2 5’UTR-R), the ORF of the *Hph* gene (LpBBS2 Hph-F and LpBBS2 Hph-R), and part of the *LpBBS2* ORF (1271-1870, LpBBS2 3’ORF-F and LpBBS2 3’ORF-R). The fusion PCR product was cloned into the EcoRV site of pBluescript II SK(+), and the linearized plasmid DNA (10 μg) with XhoI was used for electroporation of *L. passim* expressing both Cas9 and *LpBBS2* gRNA, as described above.

After electroporation, *L. passim* resistant to blasticidin, G418, and hygromycin were selected, and single parasites were cloned by serial dilutions in a 96-well plate. The genotype of each clone was initially determined through the detection of 5’ wild-type (WT) and knock-out (KO) alleles for *LpBBS2* by PCR. After identifying heterozygous (+/-) and homozygous (-/-) KO clones, their 5’WT (LpBBS2 5’UTR-Up and LpBBS2-47R), 5’KO (LpBBS2 5’UTR-Up and Hyg-159R), 3’WT (LpBBS2-1216F and LpBBS2-1910R), and 3’KO (Hyg-846F and LpBBS2-1910R) alleles were confirmed by PCR using specific primer sets.

### RT-PCR

Total RNA was extracted from WT, *LpBBS2* heterozygous, and homozygous mutant parasites using TRIzol reagent (Sigma-Aldrich). Reverse transcription of 0.2 μg of total RNA was performed using ReverTra Ace (TOYOBO) and random primers, followed by PCR with GoTaq Green Master Mix (Promega). We detected *LpBBS2* mRNA by nested PCR by running the 1st PCR with LpSL-F-1st and LpBBS2-70R primers, followed by the 2nd PCR with LpSL-F-2nd and LpBBS2-47R primers. To detect *LpGAPDH* mRNA, we used LpSL-F and LpGAPDH-R primers.

### Culture, flagellar and cell body length measurement, and motility measurement of *L. passim*

For culture, flagellar and cell body length measurement, and motility measurement, we inoculated WT and *LpBBS2*-deficient parasites into the culture medium at 10⁴ cells/mL at 30 °C. The number of parasites was counted daily using a hemocytometer, and images of the cultured parasites were captured simultaneously for 5 days. We measured the length of both the flagellum and the cell body of individual parasites at 3 days after culture using phase images and Image-J. To record the movement of parasites, videos were created by taking images for 1 minute every second. The movement of parasites was tracked using TrackMate v7.10.2 (Ershov, Phan et al. 2022) as a Fiji (Schindelin, Arganda-Carreras et al. 2012) plugin. The videos were imported to Fiji, converted to 8-bit grayscale, and the brightness and contrast were adjusted for better tracking. The Laplacian of Gaussian detector was used with an estimated object diameter of 30.0-36.0 pixels and a quality threshold of 0.2-0.5 for the detection of individual parasites. The Simple Linear Assignment Problem tracker was used to track parasites by adjusting the linking maximum distance and gap-closing maximum distance to 35.0-200.0 pixels, and the gap-closing maximum frame gap to 1. To examine the growth rates of WT and *LpBBS2*-deficient parasites at 21 °C, the cells were inoculated at 10^5^ cells/mL. We characterized the phenotypes of two independent *LpBBS2*-deficient clones (D8 and G1), and their phenotypes were the same. Therefore, we used the D8 clone for further experiments.

### Honey bee infection

To infect honey bees with *L. passim*, parasites collected during the logarithmic growth phase (5 × 10⁵ cells/mL) were washed with PBS and suspended in sterile 10% sucrose/PBS at 5 × 10⁴ cells/μL. Newly emerged honey bee workers were collected by placing frames with late pupae in a 33 °C incubator and were starved for 2-3 hours. Twenty individual honey bees were fed 2 μL of the sucrose/PBS solution containing either WT or *LpBBS2*-deficient parasites (10⁵ cells in total). The infected honey bees were maintained in metal cages at 33 °C for 14 days and then frozen at –80 °C. This experiment was repeated three times. We sampled eight honey bees from each of the three experiments, thus analyzing 24 honey bees in total infected with either WT or *LpBBS2* mutants. Genomic DNA was extracted from the whole abdomens of individual bees using DNAzol reagent. We quantified *L. passim* in the infected honey bees by qPCR using LpITS2-F and LpITS2-R primers, which correspond to part of the internal transcribed spacer region 2 (ITS2) in the ribosomal RNA gene. Honey bee *AmHsTRPA* was used as the internal reference with AmHsTRPA-F and AmHsTRPA-R primers (Liu, Lei et al. 2020). The relative abundance of *L. passim* in individual honey bees (24 each infected by WT or mutant *L. passim*) was calculated using the ΔCt method, setting one sample infected by WT as 1. Statistical analysis was performed using the Brunner-Munzel test. All the primers used are listed in Supplementary file 2.

### Localization of FCaBP1N16-GFP and FCaBP2N16-GFP and their association with lipid rafts in WT and *LpBBS2*-deficient *L. passim*

We digested plasmid DNA encoding either FCaBP1N16-GFP or FCaBP2N16-GFP with BamHI and XhoI, followed by subcloning into the same sites of pTrex-n-eGFP plasmid DNA, wherein the neomycin resistance gene was replaced by the bleomycin resistance gene. WT and *LpBBS2*-deficient *L. passim* were electroporated with the plasmid DNA and selected by either zeocin (for WT) or hygromycin and zeocin (for *LpBBS2* mutants). The GFP and DIC images of parasites were captured as described above. Using Image-J, we measured the mean fluorescence of FCaBP1N16-GFP in both the entire flagellum and cell body of individual parasites, as well as the background. We then calculated the ratio between the fluorescence in the flagellum and the cell body after subtracting the background fluorescence. Statistical comparison was performed using the Brunner-Munzel test.

To examine the association of FCaBP1N16-GFP and FCaBP2N16-GFP with lipid rafts in WT and *LpBBS2* mutant parasites, we collected the cells in the mid-log growth phase, washed them twice in 0.5 mL PBS, and resuspended them at 10⁷ cells/mL in 150 μL of 1% Triton X-100 in PBS, pre-equilibrated to either 4, 20, or 37 °C. The cells were incubated at the same temperature for 10 minutes and centrifuged at 1000 × g for 5 minutes. The supernatants were removed from the pellets and transferred to new tubes, and the pellets (insoluble fraction) were resuspended in 150 μL of 1% Triton X-100 in PBS. The supernatants were centrifuged again at 17,000 × g for 10 minutes at either 4, 20, or 37 °C, and the supernatants (soluble fraction) were collected. Both fractions were mixed with 50 μL of 4 × SDS-PAGE sample buffer, heated at 95 °C for 5 minutes, and 35 μL of each sample was applied to 12% SDS-PAGE. The proteins were detected by western blot using anti-GFP antibody as described above. We used Image-J to measure the band intensities of FCaBP1N16-GFP and FCaBP2N16-GFP in the pellet and soluble fractions after background subtraction. Then, we calculated the ratio of pellet to soluble fraction and used Welch’s t-test for statistical comparison.

### Transcriptome Analysis of WT and LpBBS2-Deficient *L. passim*

We cultured WT and LpBBS2-deficient parasites (clones D8 and G1) in three 10 cm culture plates for each genotype at 30°C, and collected the cells during the mid-log growth phase. Total RNA was independently extracted from each plate and analyzed by RNA-seq. All samples were sequenced on the DNBSEQ-T7 platform at the Beijing Genomics Institute, yielding at least 6 GB of clean data per sample.

Since the genome sequence of *L. passim* (strain 422, GenBank assembly: GCA_034478905.1) in NCBI lacks gene annotation information, we first annotated *L. passim* protein-coding genes. Gene annotation for the *L. passim* genome sequence was conducted using the BRAKER pipeline (Hoff, Lange et al. 2016) with *Crithidia bombi*, *Leptomonas pyrrhocoris*, and *Leptomonas seymouri* as reference organisms. The gffread program (http://ccb.jhu.edu/software/stringtie/gff.shtml) was used to convert the *L. passim* GFF to a GTF file, and we generated the genomic indices from the converted GTF file using genomeGenerate from STAR (Version: 2.7.0) (Dobin, Davis et al. 2013)

The quality of all RNA-seq data was analyzed using FastQC (Brown, Pirrung et al. 2017) and pruned using SOAPnuke (version 2.2.1). After trimming low-quality reads and removing reads with more than 5% adapter and N bases, more than 92% of the reads in each sample were retained for analysis. The clean RNA-seq reads were then mapped to the STAR index. The assembled reads were aligned to the *L. passim* genome using alignReads in STAR, and coordinate-sorted BAM files were generated. The mapping rates of the nine samples to the *L. passim* genome ranged from 73.05 % to 73.78 %. Principal component analysis (PCA) of the nine RNA-seq samples was conducted using the gmodels package (version 2.18.1) in R. We used htseq-count (Anders, Pyl et al. 2015), developed with HTSeq, to count the overlap of reads mapped to the GFF features as a preprocessing step for differential gene expression analysis. Raw read counts were analyzed in RStudio (Version 1.2.1335) using the linear contrast function in the DESeq2 package (Version 1.24.0) from Bioconductor (Robinson, McCarthy et al. 2010). Normalization was performed using shrinkage estimation in DESeq2 (Love, Huber et al. 2014). Reads with fewer than one count per million mapped reads were removed from the analysis. The Benjamini-Hochberg method was applied to control the false discovery rate (FDR) and log2 fold change (FC) across the detected loci. Differentially expressed genes (DEGs) between two samples (WT vs. D8, WT vs. G1, and D8 vs. G1) were identified using a threshold of FDR < 0.05 and log2 FC > 1. For scatter plot analysis, the logarithmic values of gene expression levels of two samples are plotted on x– and y-axis, respectively.

GO enrichment analysis of DEGs was performed using the clusterProfiler package (version 3.18.0) (Yu, Wang et al. 2012), and all statistical analyses were conducted in RStudio (Version 1.2.1335). After obtaining the enriched GO terms, we identified the most specific ones using a cutoff value of FDR < 0.05. The accession numbers for the RNA-seq data are SAMN44080493, SAMN44080494, SAMN44080495, SAMN44080496, SAMN44080497, SAMN44080498, SAMN44080499, SAMN44080500, and SAMN44080501.

## Author contributions

TK conceived and designed research strategy and wrote the paper. XY performed the part of experiments. XY and TK analyzed data.

## Acknowledgements

We thank Shangqing Li and Carmen Lau for their contribution to this study.

## Data availability statement

All data are included as tables, figures and Supplementary Information in the article. The RNA-seq data were deposited to NCBI with the accession numbers listed in the article.

---

**Supplementary Table 1.** Normalized read counts of dynein heavy and light chain mRNAs in wild-type and *LpBBS2* mutants.

| Gene | Log2 fold<br>change | Adjusted *P*-<br>value | WT_1 | WT_2 | WT_3 | D8_1 | D8_2 | D8_3 |
| --- | --- | --- | --- | --- | --- | --- | --- | --- |
| Dynein heavy chain | 0.190670529 | 0.003065021 | 1476.723 | 1414.448 | 1435.998 | 1764.998 | 1646.982 | 1529.304 |
| Dynein light chain | 0.25046061 | 9.23708E-06 | 1545.993 | 1578.368 | 1602.009 | 1776.286 | 1894.122 | 1949.585 |

| Gene | Log2 fold<br>change | Adjusted *P*-<br>value | WT_1 | WT_2 | WT_3 | G1_1 | G1_2 | G1_3 |
| --- | --- | --- | --- | --- | --- | --- | --- | --- |
| Dynein heavy chain | 0.310218335 | 2.59169E-06 | 1494.987 | 1433.25 | 1453.557 | 1949.186 | 1668.647 | 1814.946 |
| Dynein light chain | 0.251208148 | 6.86015E-05 | 1565.114 | 1599.349 | 1621.598 | 2047.353 | 1834 | 1814.946 |

**Supplementary Figure 1.**
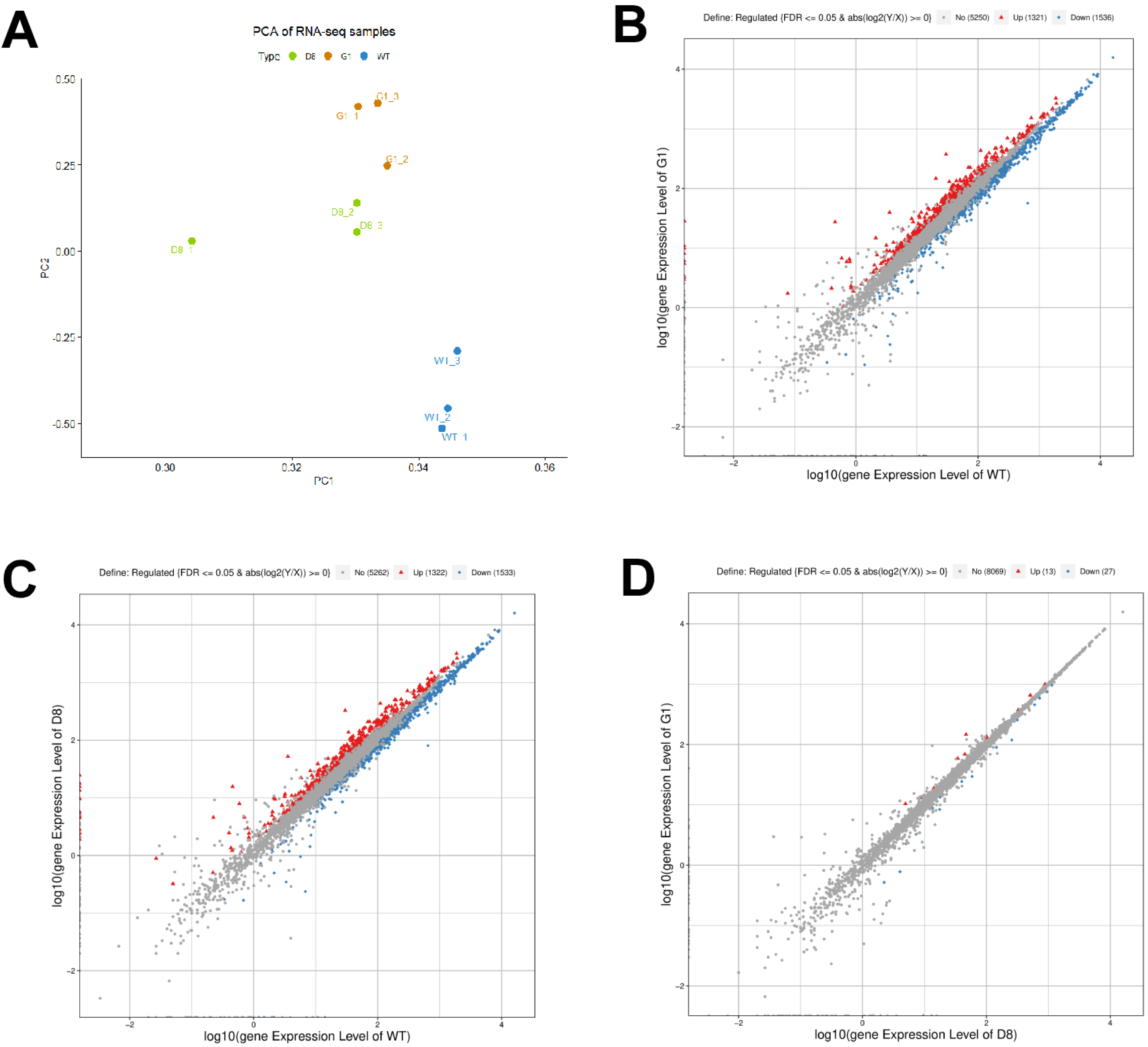
Comparison of RNA-seq data for WT and LpBBS2-deficient parasites. (A) Principal component analysis of RNA-seq samples from WT (blue), *LpBBS2* mutant clone D8 (green), and clone G1 (beige). (B-D) Scatter plot analysis showing upregulated (red), downregulated (blue), and non-significantly different genes between WT and clone G1 (B), WT and clone D8 (C), or clone D8 and clone G1 (D).

**Supplementary video 1. Movement of wild-type *L. passim***

**Supplementary video 2. Movement of LpBBS2-deficient *L. passim***

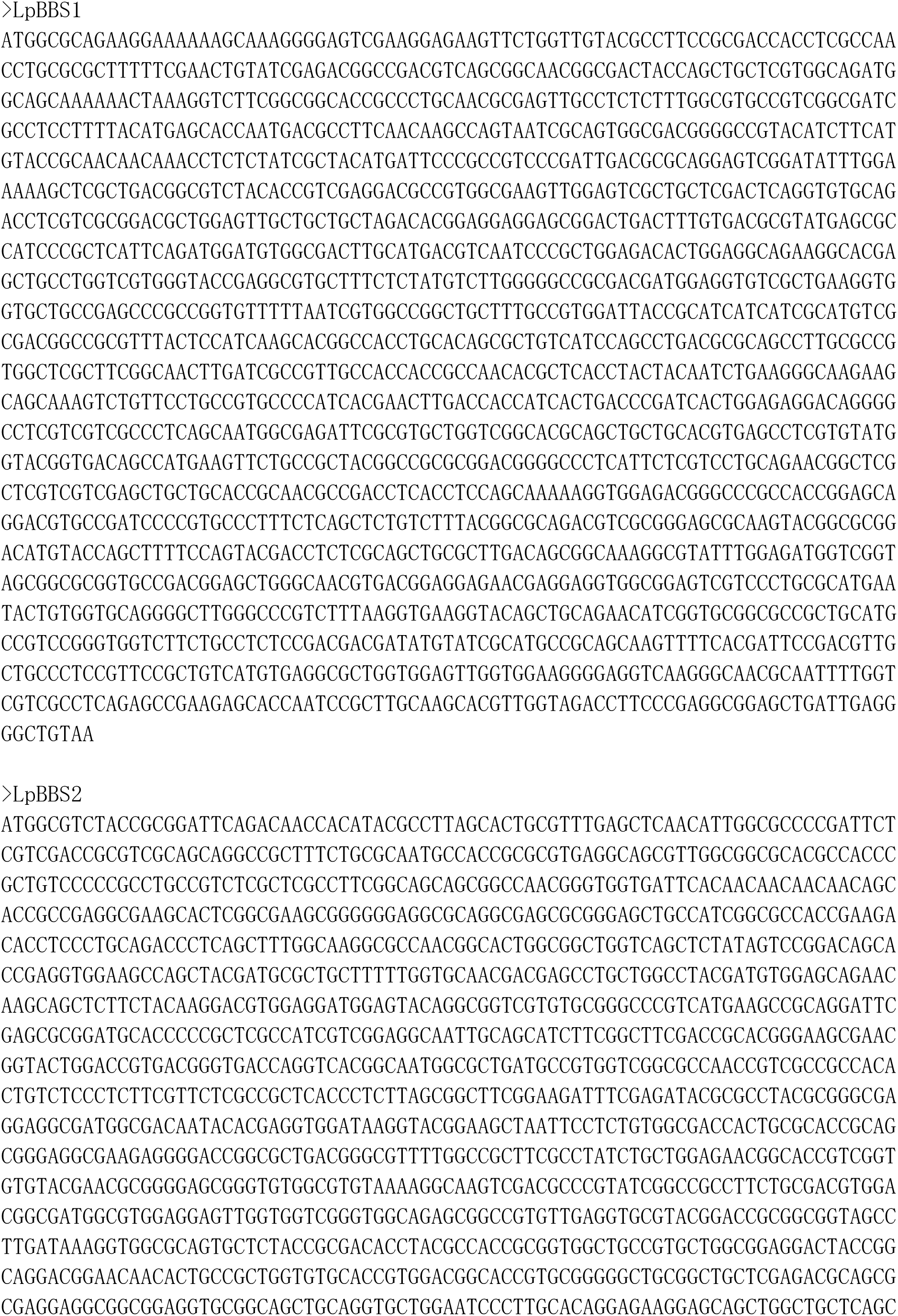
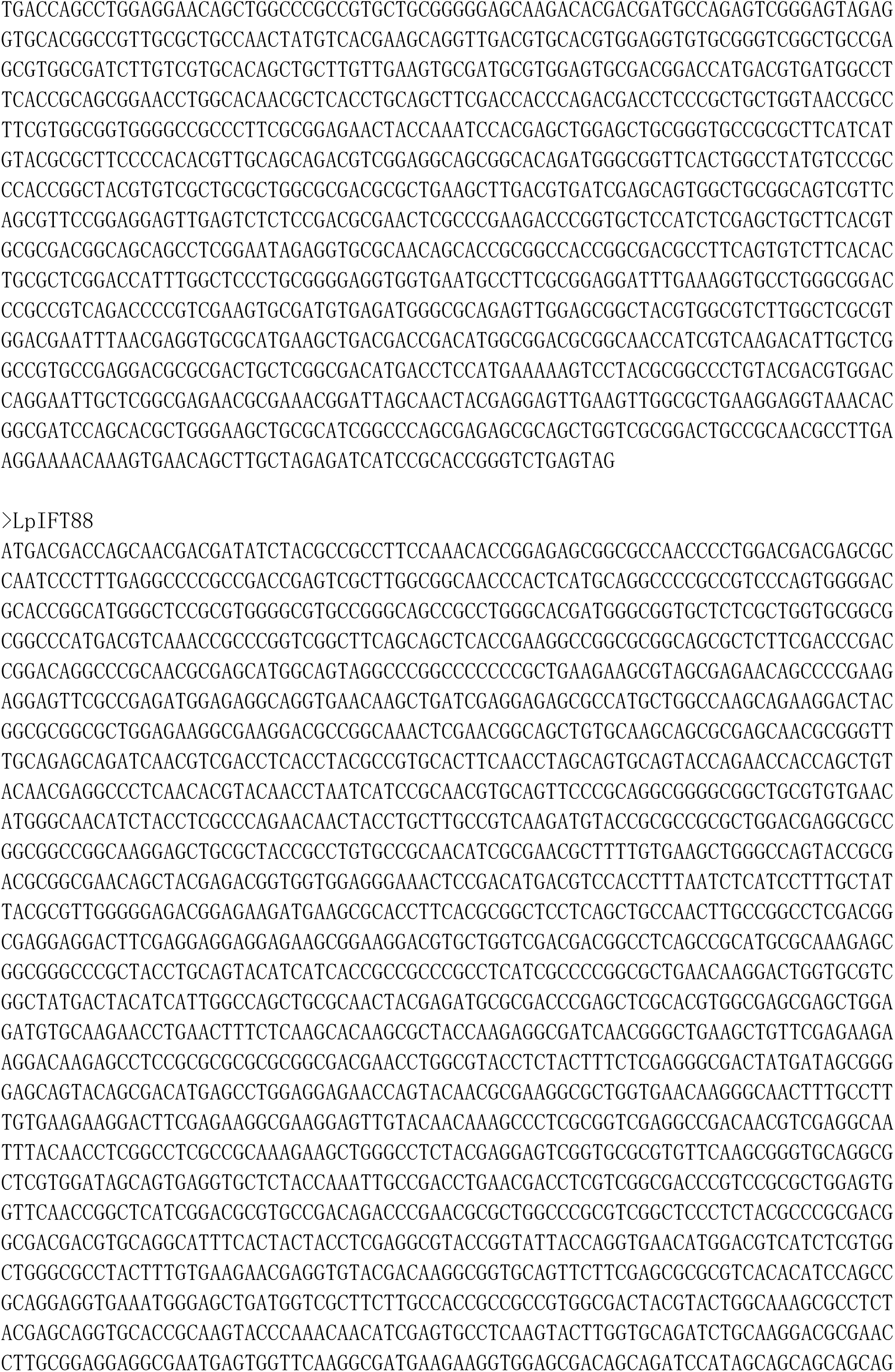
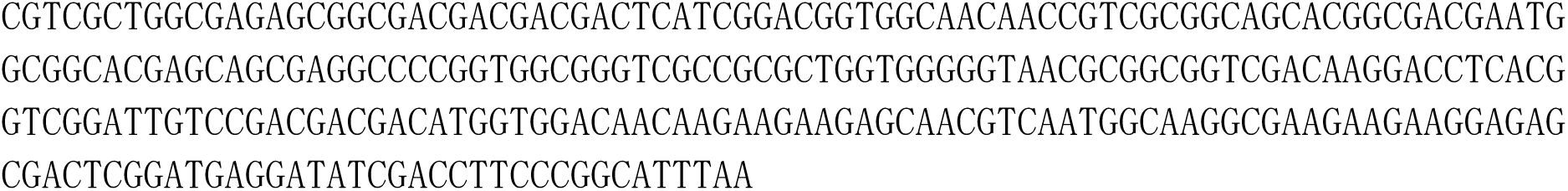
Supplementary file 1.

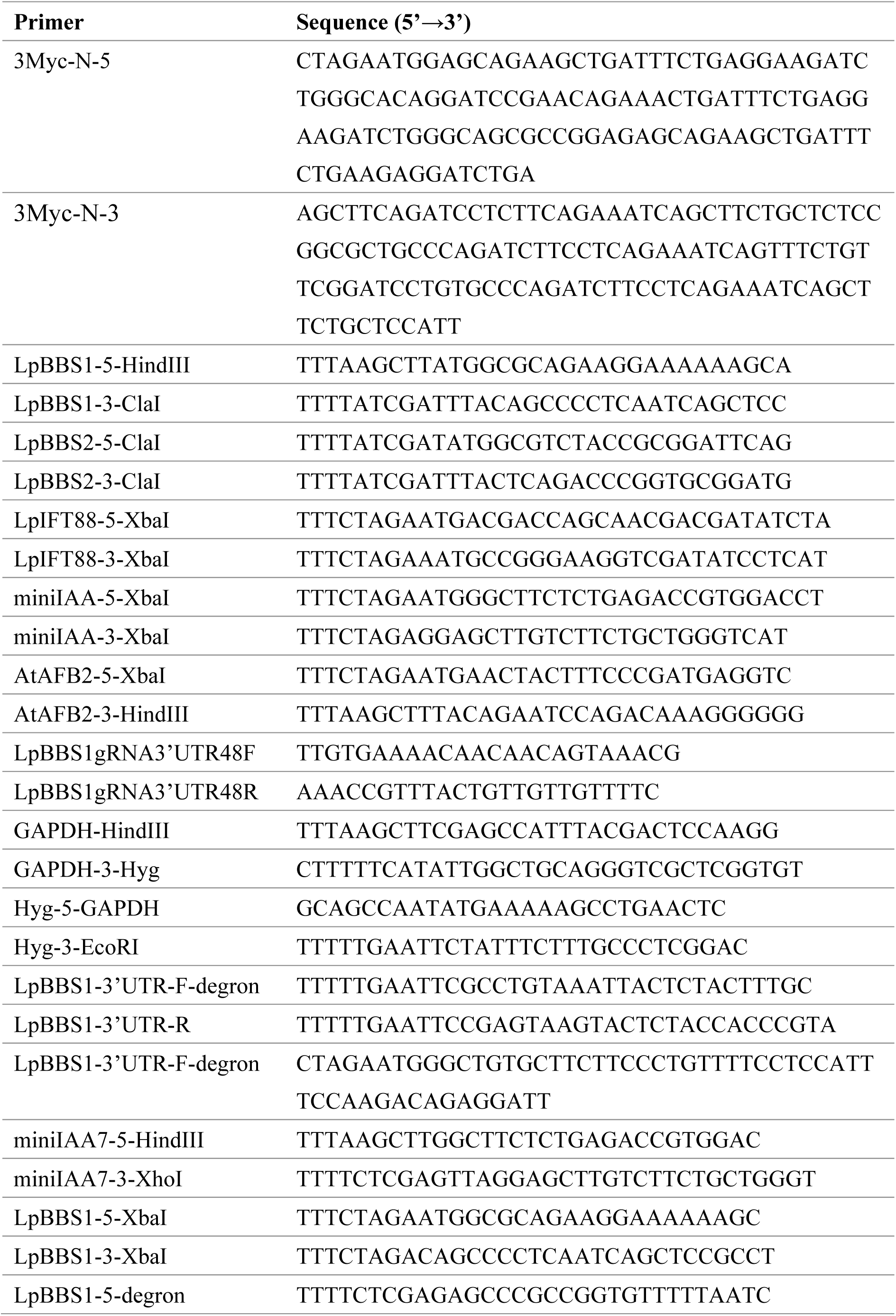
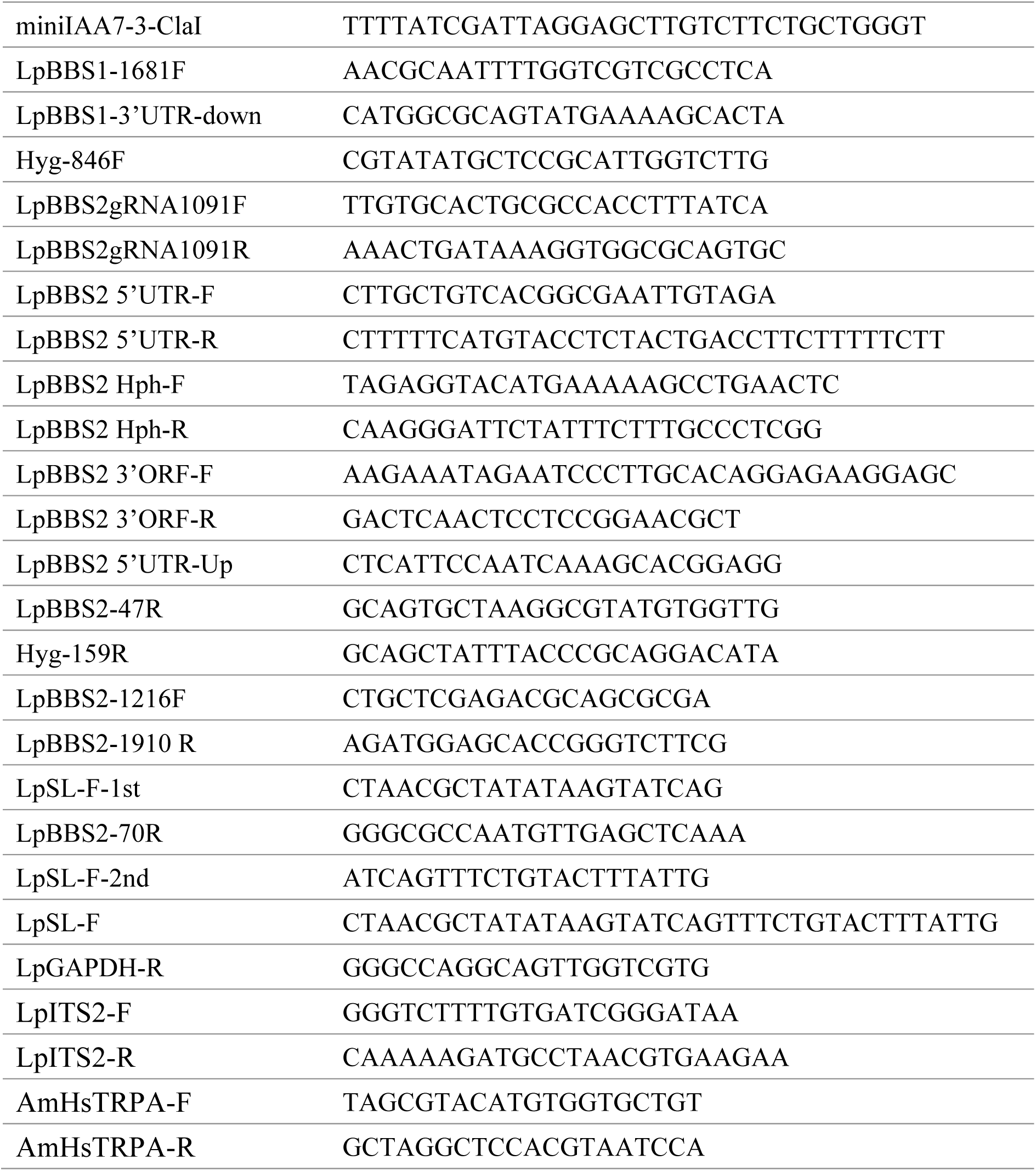
Supplementary file 2.

